# Network Signatures of Disease Progression and Core Symptoms in Dementia with Lewy Bodies Distinct from Alzheimer’s Disease

**DOI:** 10.1101/2025.07.17.665279

**Authors:** Vincent Gabriel, Elena Chabran, Marion Sourty, Benjamin Cretin, Nathalie Philippi, Candice Muller, Pierre Anthony, Catherine Demuynck, Paulo Loureiro de Sousa, Anne Botzung, Léa Sanna, Olivier Bousiges, Frédéric Blanc

## Abstract

**Background:** Resting-state fMRI studies in dementia with Lewy bodies (DLB) and Alzheimer’s disease (AD) have described connectivity alterations in large-scale brain networks. However, little is known about functional changes across disease stages, particularly in DLB.

**Objective:** To investigate functional connectivity of key brain networks in DLB patients at different stages, compare them to AD and healthy controls (HC) and examine associations with core clinical symptoms.

**Methods:** Ninety DLB patients, comprising 63 with mild cognitive impairment (MCI-DLB) and 27 with dementia (d-DLB), along with 25 AD patients (11 MCI-AD and 14 d-AD) and 34 HC underwent clinical, neuropsychological and resting-state fMRI assessment. ROI-to-ROI analyses were performed using the CONN toolbox, (*p*_FDR_<0.05).

**Results:** The DLB group showed reduced functional connectivity within the salience network (SN) compared with HC, but did not differ from AD. While MCI-DLB patients showed no significant differences, d-DLB patients showed reduced SN and frontoparietal network (FPN) connectivity compared to HC and AD. SN connectivity was associated with severity of fluctuations and FPN connectivity with REM sleep behavior disorder and cognitive decline in DLB. In contrast, in the AD group, decreased default mode network (DMN) connectivity was associated with lower MMSE scores.

**Conclusion:** SN and FPN connectivity impairments relate to disease progression and core clinical features in DLB, whereas DMN connectivity is linked to cognitive decline in AD. These distinct patterns highlight divergent paths of network dysfunction in the two diseases, offering insight into their underlying mechanisms and clinical expression.

## Introduction

Dementia with Lewy bodies (DLB) is the second most common progressive cognitive disorder after Alzheimer’s disease (AD), accounting for approximately 15-25% of neuropathologically confirmed cases.^1^ Clinically, DLB presents a complex profile that overlaps with both AD and Parkinson’s disease, including cognitive decline, fluctuations in cognition and attention, recurrent visual hallucinations, parkinsonism, and REM sleep behavior disorder (RBD).^1^ These overlapping symptoms complicate the differential diagnosis, especially in the early stages of the disease, where differentiating mild cognitive impairment (MCI) from advanced dementia is critical.^2^

Previous studies using resting-state functional magnetic resonance imaging (fMRI) have shown functional connectivity alterations within key brain networks in DLB, such as the default mode network (DMN),^3,4,5^ the salience network (SN),^4,6^ the frontoparietal network (FPN),^4,7,8^ and different motor networks.^5,7,9^ These networks support cognitive functions, attention and executive processes, which are progressively impaired in DLB patients. Although some studies have compared connectivity patterns across DLB, AD and healthy controls, our knowledge of how these connectivity changes evolve as the disease progresses from the MCI stage to more severe dementia remains limited. Moreover, few studies have directly examined how these network alterations relate to clinical symptoms.

The SN, mainly involving the anterior insula and anterior cingulate cortex, plays a critical role in detecting salient stimuli and regulating attention and emotional responses.^10^ Its connectivity impairments have been consistently reported in neurodegenerative diseases, including DLB, and appear to contribute to the attentional and psychiatric symptoms.^6,11^ The FPN, involving structures such as the lateral prefrontal cortex (PFC) and the posterior parietal cortex (PPC),^12^ is implicated in executive control regulation and cognitive flexibility.^13,14^ Dysfunctions in the FPN are associated with cognitive decline in MCI patients.^15^ Finally, the DMN, mainly involving the posterior cingulate cortex (PCC) and medial PFC, supports self-referential thoughts, memory consolidation, and future planning.^16^ In AD, DMN connectivity appears to be significantly reduced, particularly in the posterior regions, correlating with memory impairments.^17,18,19,20^ In contrast, DLB patients tend to show more subtle changes in DMN connectivity.^5,6,9,21^ How these network alterations relate to dementia progression remains poorly understood, particularly in DLB.

The MCI stage is characterized by cognitive decline that is noticeable but not yet severe enough to compromise the patient’s independence.^22^ To date, only one study has specifically investigated whole-brain connectivity in MCI DLB patients and failed to detect significant differences compared to healthy controls.^23^ More focused analyses of network-specific changes could serve as reliable biomarkers of dementia progression in DLB.

This study investigated both between- and within-network functional connectivity changes in a large cohort of DLB patients, at both the prodromal (MCI-DLB) and demented (d-DLB) stages, compared to AD (MCI-AD and d-AD) patients and healthy controls (HC). We hypothesized that core networks involved in DLB, such as the SN, would show progressively impaired connectivity as the disease advances towards dementia. These changes could not only distinguish DLB from HC, but also from AD. Furthermore, we hypothesized that these alterations would correlate with key clinical features of DLB. These relationships could improve our understanding of the neural mechanisms underlying cognitive decline in DLB, enhance diagnostic accuracy, and help identify potential targets for neuromodulation-based therapeutic interventions in the long term.

## Methods

### Participants

Participants were recruited among the AlphaLewyMA longitudinal research protocol (ClinicalTrials.gov, registration number: NCT01876459; URL: https://clinicaltrials.gov/ct2/show/NCT01876459) initiated in 2013 at the University Hospital of Strasbourg’s tertiary Memory Clinic (CM2R), France. Patients underwent detailed clinical and neuropsychological evaluations (Table 1), blood sampling, MRI acquisitions, and lumbar puncture. DLB patients were diagnosed according to the DLB consensus criteria,^1^ which include cognitive and attentional fluctuations,^24^ visual hallucinations,^25^ parkinsonism features^26^ and RBD.^27^ AD patients were selected according to Albert’s and Dubois’ criteria.^28,29^ Patients were further categorized into prodromal (MCI-DLB, MCI-AD) and demented (d-DLB, d-AD) subgroups based on their Instrumental Activities of Daily Living (IADL) score and McKeith’s criteria.^2^ Patients with missing or poor-quality imaging data were excluded from the study. A total of 90 DLB patients (63 MCI-DLB and 27 d-DLB), 25 AD patients (11 MCI-AD and 14 d-AD) and 34 HC matched for age and gender were included for these analyses (Table 1).

**Table 1.** Sociodemographic and Clinical Characteristics of the DLB, AD, and HC groups.

| Characteristic |  | DLB<br>*n* = 90 | AD<br>*n* = 25 | HC<br>*n* = 34 | Test statistics <sup>d</sup>,<br>*p* | Post-hoc<br>analyses |
| --- | --- | --- | --- | --- | --- | --- |
| **Age (years)** |  | 69.5 ± 9.5 | 73.2 ± 8.2 | 68.1 ± 8.1 | *F* = 2.394<br>*p* = 0.095 | / |
| **Gender (F/M) <sup>a</sup>** | | 53/37 | 15/10 | 21/13 | $\chi^2$ = 0.040<br>*p* = 0.980 | / |
| **MMSE score** |  | 25.7 ± 3.8 | 24.0 ± 3.2 | 28.9 ± 1.0 | *H* = 42.27<br>*p* < 0.001*** | HC > DLB, AD |
| **Socio-educational level <sup>b</sup>** |  | 11.7 ± 4.2 | 11.8 ± 3.1 | 13.2 ± 2.8 | *H* = 4.12<br>*p* = 0.127 | / |
| **IADL** |  | 3.4 ± 1.0 | 3.2 ± 0.9 | 4.0 ± 0.0<br>(1 ND) | *H* = 16.78<br>*p* < 0.001*** | HC > DLB, AD |
| **Fluctuations** |  | 1.9 ± 1.2 | 0.5 ± 0.8 | 0.3 ± 0.5 | *H* = 56.02<br>*p* < 0.001*** | DLB > HC, AD |
| **Hallucinations** |  | 1.7 ± 1.9 | 0.2 ± 0.4 | 0.2 ± 0.4 | *H* = 35.43<br>*p* < 0.001*** | DLB > HC, AD |
| **Parkinsonism** | **Akinesia** | 0.8 ± 0.7 | 0.0 ± 0.2 | 0.0 ± 0.2 | *H* = 53.06<br>*p* < 0.001*** | DLB > HC, AD |
|  | **Rigidity** | 0.9 ± 0.7 | 0.0 ± 0.2 | 0.0 ± 0.2 | *H* = 54.45<br>*p* < 0.001*** | DLB > HC, AD |
| | **Tremors at rest <sup>c</sup>** | 28.89% | 4.00 % | 5.88% | $\chi^2$ = 12.78<br>*p* < 0.01** | DLB > HC, AD |
| **RBD** |  | 1.1 ± 0.9<br>(2 ND) | 0.2 ± 0.5 | 0.4 ± 0.6 | *H* = 31.81<br>*p* < 0.001*** | DLB > HC, AD |

All values are presented as means ± standard deviation unless otherwise stated.
<sup>a</sup> Sex-ratio (Female/Male). <sup>b</sup> Number of years of education. <sup>c</sup> Percentage of patients with deficits. <sup>d</sup> Statistical analysis performed using the one-way ANOVA (*F*) with Tukey's multiple comparison test; the Kruskal-Wallis with Dunn's multiple comparison post-hoc test (*H*); or the chi-square test ( $\chi^2$ ) with Bonferroni correction for multiple comparisons.
DLB, dementia with Lewy bodies; AD, Alzheimer's disease; HC, healthy controls; MMSE, Mini-Mental State Examination; IADL, Instrumental Activities of Daily Living; RBD, REM sleep behavior disorder.

All participants provided written informed consent in accordance with the Declaration of Helsinki and the study was approved by the local ethics committee of Eastern France (IV).

### fMRI Data Acquisition

Functional and anatomical images were acquired using a 3 Tesla MR scanner (Verio 32-channel Tim Siemens scanner; Siemens, Erlangen, Germany). Resting-state fMRI (BOLD) and arterial spin-labeling (ASL) sequences were used to capture 121 whole-brain T2*-weighted echo planar images (repetition time = 3s; flip angle = 90°; echo time = 21ms; resolution = 38×64×28; field of view = 152×256×112 mm^2^; 4mm isotropic voxels). The first volume, acquired for ASL assessment, was excluded from the connectivity analysis. Additionally, a whole-brain T1-weighted anatomical image was acquired using a volumetric magnetization-prepared rapid acquisition with gradient-echo (MPRAGE) sequence (repetition time = 1900ms; flip angle = 9°; echo time = 2.52ms; field of view = 256×256mm^2^; resolution 256×256×256 voxels; slice thickness = 1mm).

### fMRI Data Preprocessing

Functional images were pre-processed using the Statistical Parametric 12 package (SPM12, The Wellcome Trust Centre for Neuroimaging, London, UK) running on Matlab R2023a (MathWorks, Natick, MA, USA). The preprocessing steps included: low-pass filtering of ASL frequencies at 0.112 Hz (based on ^30^); slice-timing correction; motion and B_0_ field corrections; correction of excessive motion or intensity fluctuations using ArtRepair; co-registration of the fMRI images to the T1-weighted anatomical images; spatial normalization to the Montreal Neurological Institute space using the DARTEL algorithm, followed by smoothing with an 8mm Gaussian kernel; and denoising via the CONN toolbox.^31^

### Clinical Characteristics Analysis

The clinical and sociodemographic characteristics of the patients among the three groups were compared using JASP software (http://jasp-stats.org). After testing for normality and variance homogeneity, between-group differences were evaluated using either a one-way analysis of variance (ANOVA) with Tukey’s post-hoc test, or a Kruskal-Wallis test, followed by Dunn’s multiple comparison test for post-hoc analyses. Categorical data were analyzed with a χ^2^ test with Bonferroni correction for multiple comparisons. Statistical significance was set at *p* < 0.05.

### Resting-state Functional Connectivity Analysis

#### ROI-to-ROI Analysis

Seed-based functional connectivity analyses were performed using the CONN toolbox, running on Matlab R2023a. Thirty-two regions of interest (ROIs) corresponding to the main nodes of the DMN, FPN, SN, dorsal attention network (DAN), language network (LN), sensorimotor network (SMN), visual network (VN), and cerebellar network (CN) were selected from the Harvard-Oxford ROI atlas from the CONN toolbox.^32^

Individual ROI-to-ROI functional connectivity matrices were created using Pearson’s correlation between BOLD time series of each pair of ROIs. The six motion parameters resulting from preprocessing were included as covariates of non-interest. Correlation values were converted to z-scores via Fisher’s transformation to meet the normality required for second-level analysis. These matrices were then entered into a general linear model corrected for age, gender and total intracranial volume (TIV) for group comparisons. Results were considered significant at a false discovery rate (FDR)-corrected threshold of *p*_*FDR*_ < 0.05 at the cluster level (MVPA omnibus test), and *p* < 0.05 uncorrected at the connection level.

#### Association between Functional Connectivity and Clinical Variables

Multiple regression analyses were conducted to analyze the relationship between resting-state functional connectivity and clinical scores across the groups. Age, gender, and TIV were included as covariates of no interest. Results were considered significant at *p*_FDR_ < 0.05 at the cluster level, and p < 0.05 uncorrected at the connection level.

## Results

### Sociodemographic and Clinical Characteristics

Table 1 summarizes the sociodemographic and clinical characteristics of the participants, divided into three groups: DLB (*n* = 90), AD (*n* = 25) and HC (*n* = 34). Age was similar across the groups, with no significant difference observed (*p* = 0.095). Similarly, gender distribution was also comparable between the three groups (*p* = 0.980) and no significant differences were observed regarding socio-educational level (*p* = 0.127).

Regarding cognitive performances, the HC group showed significantly higher Mini-Mental State Examination (MMSE) and IADL scores than both AD and DLB patients (*p <* 0.001). We observed a significantly higher severity of cognitive fluctuations, visual hallucinations, parkinsonism, and RBD in the DLB group compared to the HC and AD groups (*p* < 0.001; except for tremors at rest, where *p* < 0.01).

When subdividing patients into MCI and demented subgroups, the d-DLB subgroup showed lower MMSE and IADL scores (*p* < 0.001), and more pronounced akinesia and rigidity (*p* < 0.001) compared to the MCI-DLB subgroup. No significant differences were observed between the two subgroups for other clinical symptoms. Finally, the MCI-AD and MCI-DLB subgroups were matched for MMSE scores, as were the d-AD and d-DLB subgroups (*p* < 0.001). Further details of these additional results are summarized in Supplementary Table 1.

### Functional Connectivity Group Comparisons (ROI-to-ROI Analysis)

All significant functional connectivity results were reported with a significance threshold of *p* < 0.05 FDR-corrected at the cluster level and *p* < 0.05 uncorrected at the connection level.

Figure 1 presents the intra- and inter-network functional connectivity patterns in the DLB group. The strongest connectivity reductions were observed in inter-network connections, particularly between the DMN and the SN, and between the DMN and the DAN. In contrast, higher connectivity was found within each functional network. Additionally, a strong inter-network connectivity was observed involving the SN, FPN, DAN, and LN.

**Figure 1.**
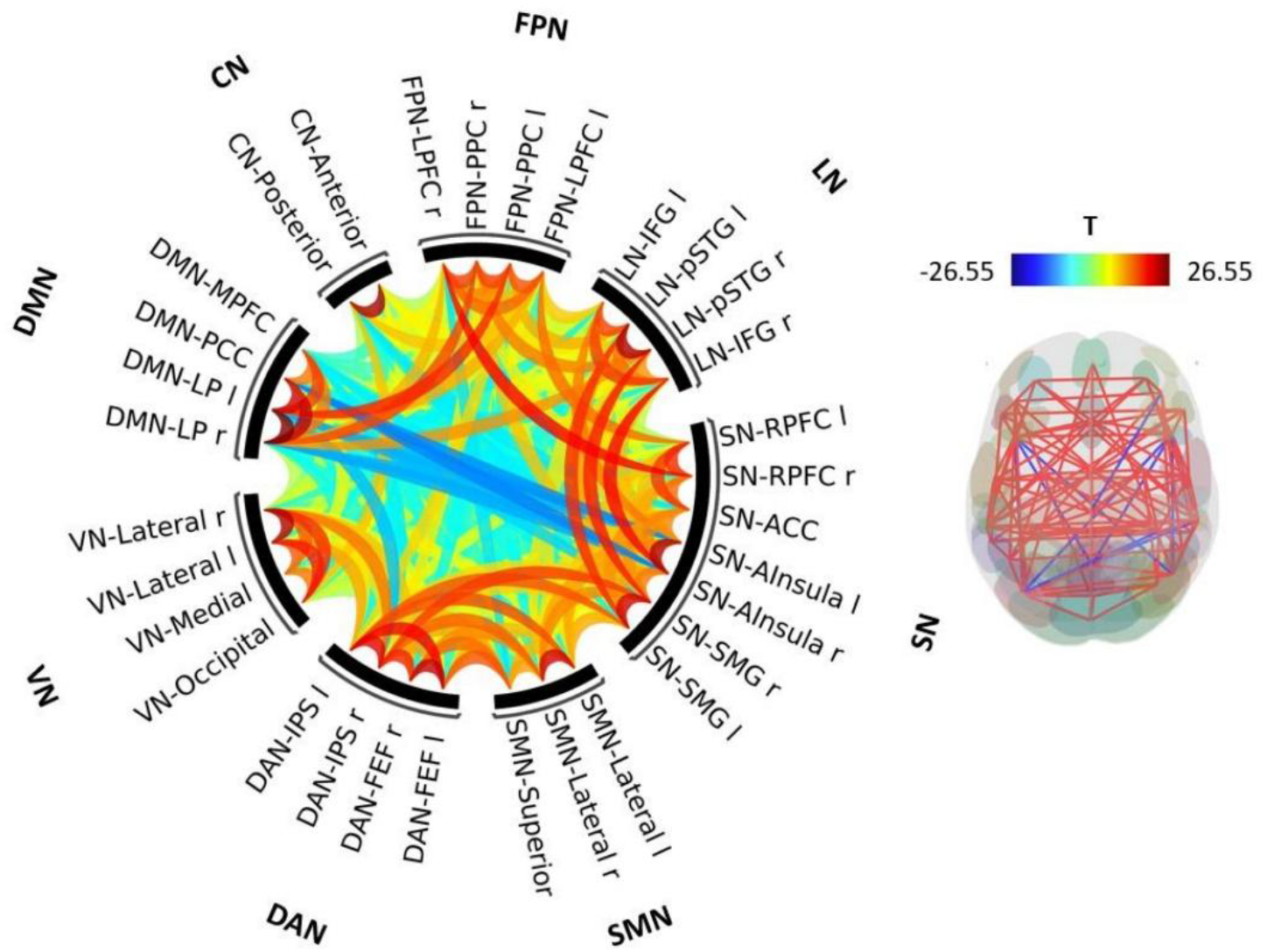
Intra- and inter-network resting-state functional connectivity in the DLB group. Significant connections were identified using a cluster threshold of p < 0.05 cluster-level pFDR corrected (MVPA omnibus test) and a connection threshold of p < 0.05 p-uncorrected. Hypoconnectivity is represented in blue, hyperconnectivity is represented in red. l, left; r, right; DMN, default mode network; CN, cerebellar network; FPN, frontoparietal network; LN, language network; SN, salience network; SMN, sensorimotor network; DAN, dorsal attention network; VN, visual network; LP, lateral parietal cortex; PCC, posterior cingulate cortex; MPFC, medial prefrontal cortex; LPFC, lateral prefrontal cortex; PPC, posterior parietal cortex; IFG, inferior frontal gyrus; pSTG, posterior superior temporal gyrus; RPFC, rostral prefrontal cortex; ACC, anterior cingulate cortex; AInsula, anterior insula; SMG, supramarginal gyrus; FEF, frontal eye fields; IPS, intraparietal sulcus.

Figure 2 illustrates the between-group differences in resting-state functional connectivity. The DLB group presented reduced connectivity within the SN compared to the HC group (Figure 2a). No significant differences were observed in the MCI-DLB subgroup. However, in the d-DLB subgroup, we observed significantly lower functional connectivity within the SN and the FPN compared to the HC group (Figure 2b).

**Figure 2.**
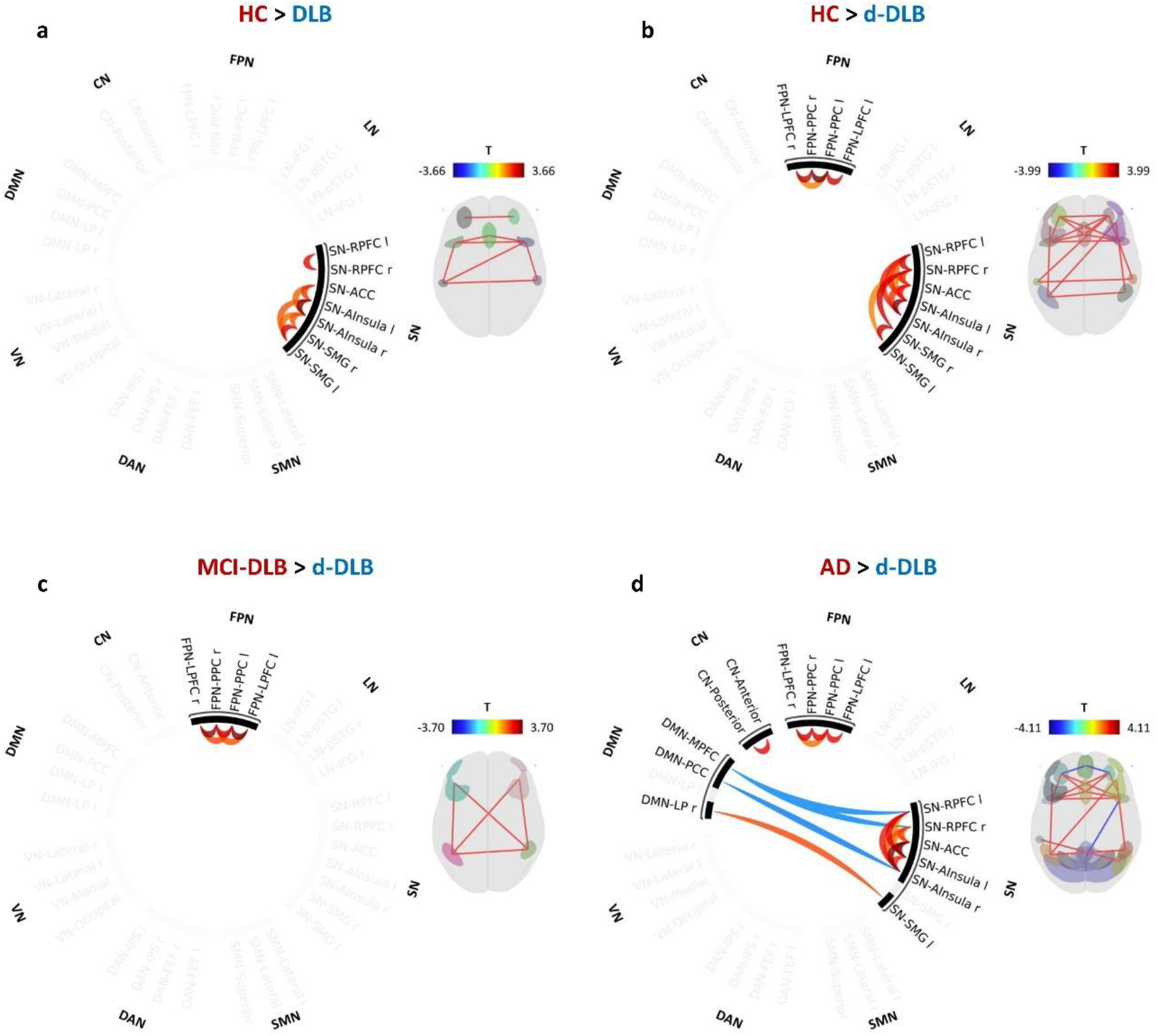
Between-group resting-state functional connectivity comparisons. This figure displays the connectivity differences between HC and DLB groups (**a**), between HC group and d-DLB subgroup (**b**), between MCI-DLB and d-DLB subgroups (**c**), and between AD group and d-DLB subgroup (**d**). Significant connections were identified using a cluster threshold of p < 0.05 cluster-level pFDR corrected (MVPA omnibus test) and a connection threshold of p < 0.05 p-uncorrected. Lower connectivity in the first group of each comparison is shown in blue; higher connectivity in the first group of each comparison is shown in red. l, left; r, right; DMN, default mode network; CN, cerebellar network; FPN, frontoparietal network; LN, language network; SN, salience network; SMN, sensorimotor network; DAN, dorsal attention network; VN, visual network; LPFC, lateral prefrontal cortex; PPC, posterior parietal cortex; RPFC, rostral prefrontal cortex; ACC, anterior cingulate cortex; AInsula, anterior insula; SMG, supramarginal gyrus; LP, lateral parietal cortex; PCC, posterior cingulate cortex; MPFC, medial prefrontal cortex.

Additionally, the comparison between the two DLB subgroups revealed significantly decreased connectivity within the FPN in the d-DLB subgroup compared to the MCI-DLB subgroup (Figure 2c).

Finally, we found no significant differences when comparing connectivity between AD and DLB patients. Again, we found no significant differences with the MCI-DLB subgroup. However, the d-DLB subgroup showed reduced connectivity within the SN, FPN, and CN compared to the AD group (Figure 2d). Additionally, the d-DLB subgroup showed reduced connectivity between the right lateral parietal cortex (DMN) and the left supramarginal gyrus (SN). In contrast, the AD group showed lower connectivity between the SN and the DMN when compared to the d-DLB subgroup, particularly between the medial PFC (DMN) and the right rostral PFC (SN), and between the PCC (DMN) and the right anterior insular cortex (SN). No significant differences were found when comparing DLB patients and their subgroups with the MCI-AD and d-AD subgroups.

### ROI-to-ROI Functional Connectivity Correlates with Clinical Symptoms

Figure 3 illustrates significant correlations between resting-state functional connectivity and clinical features in the DLB and AD groups. First, MMSE scores in DLB patients were correlated with reduced connectivity within the FPN (Figure 3a), whereas MMSE scores in AD patients were correlated with reduced connectivity within the DMN (Figure 3b).

**Figure 3.**
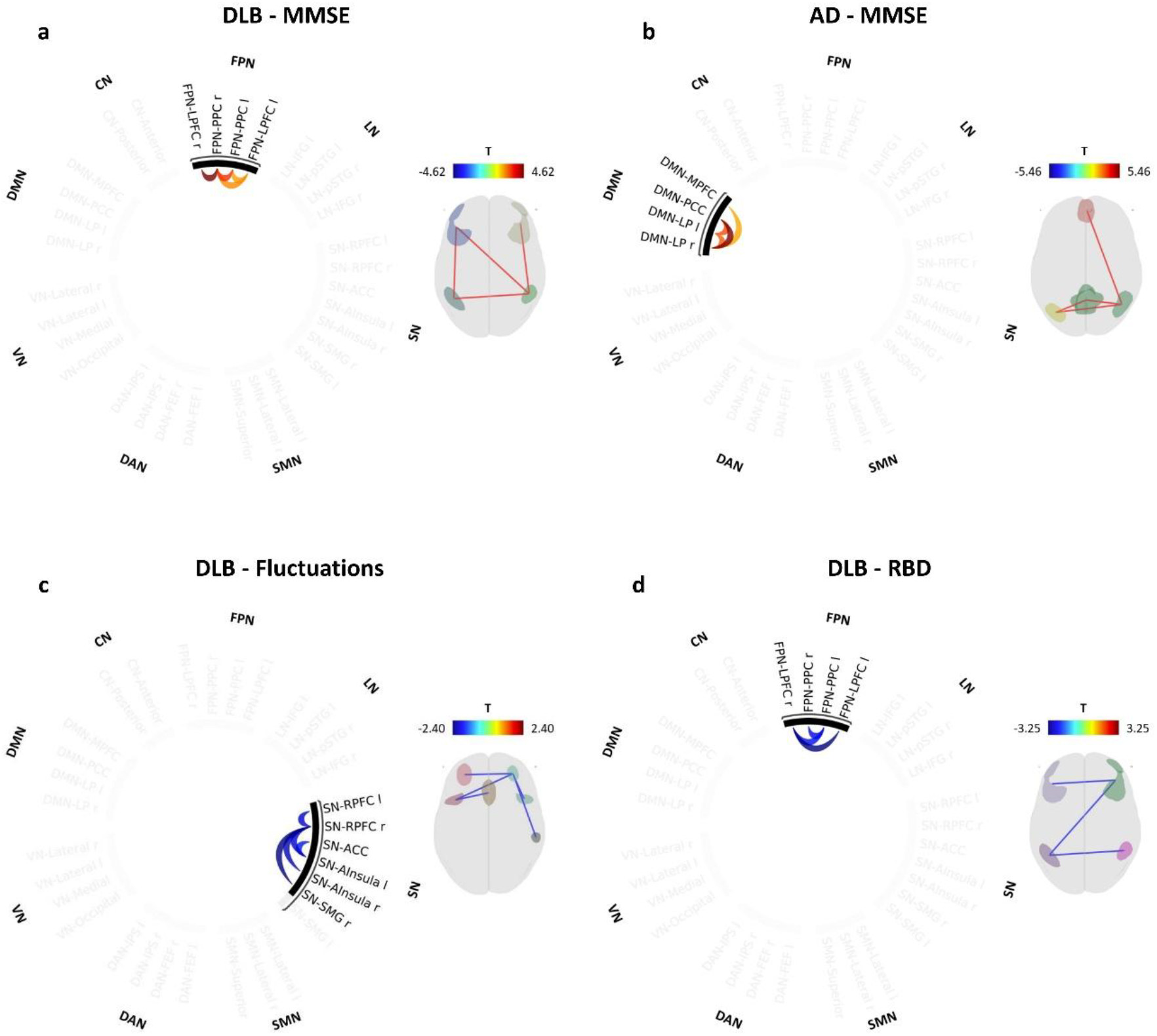
Correlations between resting-state functional connectivity and clinical features. This figure displays significant correlations between connectivity and the MMSE score in the DLB group (**a**) and the AD group (**b**), the fluctuation rate in the DLB group (**c**) and the RBD rate in the DLB group (**d**). Significant connections were identified using a cluster threshold of p < 0.05 cluster-level pFDR corrected (MVPA omnibus test) and a connection threshold of p < 0.05 p-uncorrected. Positive correlations are shown in red; negative correlations are shown in blue. DLB, dementia with Lewy bodies; AD, Alzheimer’s disease; MMSE, Mini-Mental State Examination; RBD, REM sleep behavior disorder; l, left; r, right; DMN, default mode network; CN, cerebellar network; FPN, frontoparietal network; LN, language network; SN, salience network; SMN, sensorimotor network; DAN, dorsal attention network; VN, visual network; LP, lateral parietal cortex; PCC, posterior cingulate cortex; MPFC, medial prefrontal cortex; LPFC, lateral prefrontal cortex; PPC, posterior parietal cortex; RPFC, rostral prefrontal cortex; ACC, anterior cingulate cortex; AInsula, anterior insula; SMG, supramarginal gyrus.

Additionally, in the DLB group, higher severity of fluctuations was associated with reduced connectivity within the SN (Figure 3c), while lower FPN connectivity correlated with higher RBD scores (Figure 3d). No significant correlations were found for visual hallucinations or parkinsonian symptoms.

No significant correlations between network functional connectivity and the core clinical features of DLB were observed in the AD or HC groups. When analyzing disease subgroups separately (MCI-DLB, d-DLB, MCI-AD and d-AD), no significant correlations were identified.

## Discussion

This study explored resting-state functional connectivity alterations in DLB and AD across different disease stages, compared to HC. We identified network-specific disruptions in DLB, particularly within the SN and FPN, which correlated with fluctuation severity, RBD symptoms and cognitive decline. In AD, cognitive decline was more closely related to DMN impairment. These findings highlight distinct patterns of network dysfunction in DLB and AD, and their relevance to clinical features.

### Global Functional Connectivity in DLB Patients

Our analysis of connectivity strengths across the DLB group revealed overall preserved patterns of intra-network organization, particularly within core functional networks (Figure 1). As expected, intra-network connections were generally stronger, reflecting well-known large-scale network architecture. Notably, we observed reduced connectivity between the DMN and both the SN and DAN, in line with the functional opposition between the DMN and attention-oriented networks.^33,34^ Particularly, the SN plays a switching role in response to detected salient stimuli and engages attentional systems to redirect cognitive focus.^35^ This role of the SN could also explain the strong inter-network connectivity we observe with other task-positive networks, such as the FPN or the DAN for example. These results also validate our preprocessing approach, reassuring the reliability of our data for further connectivity analyses.

### Salience Network Dysfunction in DLB

Compared to HC, DLB patients showed significant hypoconnectivity within the SN (Figure 2a), consistent with prior studies documenting dysfunctions of this network in DLB.^4,6,11^ Notably, this alteration was most evident at the dementia stage. Indeed, the MCI-DLB subgroup did not differ from HC, whereas the d-DLB subgroup had pronounced connectivity loss (Figure 2b). These results suggest that SN dysfunction emerges or intensifies as the disease progresses, in line with findings from recent studies in prodromal DLB.^23^

The SN is crucial for detecting salient stimuli and mediating switches between internal and external attention.^36^ In our cohort, reduced connectivity within the SN in DLB was significantly associated with the severity of cognitive and attentional fluctuations (Figure 3c), supporting the idea that SN integrity supports the attentional stability in these patients. This observation aligns with prior studies also linking the hallmark cognitive fluctuations with SN disruptions in DLB.^6^

Particularly, the bilateral anterior insula, a key SN hub involved in interoception, emotional regulation and consciousness,^37^ emerged as a key region affected within the SN. This region plays a central role in modulating the switch between cognitive states, which is essential for processing salient stimuli.^37,38^ Thus, it appears particularly relevant to DLB pathophysiology. Insular atrophy is frequently reported in DLB,^39^ even at the MCI stage.^40^ Notably, insular atrophy in DLB has been shown to correlate with cognitive decline.^41^ These observations suggest that structural and functional changes in the SN contribute to the attentional and cognitive instabilities that worsen with disease progression. Given that global brain atrophy is relatively mild in DLB compared to AD,^42^ such SN connectivity impairments may precede structural degeneration and help explain early clinical symptoms.

These findings highlight the critical role of SN dysfunction, particularly involving the anterior insula in maintaining attentional and cognitive stability in DLB. However, it is interesting to note that although SN impairment followed disease progression, we did not observe a direct correlation with MMSE scores. This may indicate that SN dysfunction is more closely related to the fluctuating symptomatology of the disease rather than to global cognitive decline itself.

### Frontoparietal Network Dysfunction in DLB

Changes in FPN connectivity in DLB patients followed a somewhat different trajectory. While the overall DLB group did not significantly differ from HC (Figure 2a), a significant reduction in intra-network connectivity was evident in the d-DLB subgroup (Figure 2b). These results suggest that, as for the SN, FPN alterations are linked to disease progression. However, FPN connectivity was significantly lower in the d-DLB subgroup compared to the MCI-DLB subgroup, whereas no such difference was found for SN connectivity (Figure 2c). This dissociation implies that FPN dysfunction may emerge later or evolve more sharply as dementia progresses, making it a potential marker of advanced DLB stages.

Given the FPN’s role in executive functioning and cognitive control,^12^ its disruption likely contributes to the prominent executive deficits and cognitive rigidity often observed in d-DLB.^43^ This is supported by the correlation between higher FPN connectivity and better global cognition in DLB patients (Figure 3a). These results mirror prior evidence that stronger FPN connectivity is linked to preserved key clinical features such as executive performances.^8,44^ Moreover, the fact that FPN connectivity correlated with MMSE scores, whereas SN connectivity did not, furthermore suggests that the FPN may be more specifically associated with global cognitive decline in DLB.

A novel and particularly relevant observation here is the association between reduced FPN connectivity and more severe RBD manifestations in DLB patients (Figure 3b). A possible explanation is that dysfunction in the FPN, which includes prefrontal regions involved in motor planning and inhibition,^45^ might contribute to the loss of normal REM sleep atonia and the emergence of motor behaviors during dreams.

Taken together, these findings reinforce the notion that large-scale brain network dysfunctions drive the clinical heterogeneity and disease progression in DLB.

### Comparison with Alzheimer’s Disease

In our cohort, the overall DLB group did not show significant connectivity differences from the overall AD group. However, when focusing on advanced stages, d-DLB patients exhibited more pronounced intra-network connectivity reductions in the SN, FPN and CN compared to AD patients (Figure 2d). These findings reinforce the central, and potentially DLB-specific, role of SN and FPN dysfunctions in the progression towards dementia, highlighting key differences in neuroevolutive pathways between AD and DLB.^4,5,46,47^

CN impairment, though less studied, is of interest given the cerebellum’s established involvement in motor control and balance.^48^ Cerebellar dysfunction has also been linked to visual hallucinations and shows widespread disruption in DLB.^49,50^ The reduced CN connectivity observed in d-DLB compared to AD may thus reflect the broader clinical phenotype of DLB. Although we did not find significant differences in CN connectivity between DLB patients and HC, such alterations have been recently reported.^51^

In contrast, AD patients demonstrated more prominent disruption within the DMN, with lower connectivity correlating with cognitive decline in the AD group (Figure 3b), in line with previous reports.^19^ Interestingly, the DMN-SN pathway showed a complex pattern, with both hypo- and hyperconnectivity in DLB compared to AD, depending on the regions involved. This may reflect a transitional imbalance between compensatory mechanisms and progressive disconnection. Overall, our results align with prior studies suggesting that DMN integrity is relatively preserved in DLB compared to AD,^9,52,53^ where DMN disruption is a hallmark feature,^17,54^ while DLB predominantly affects attentional and executive control networks.^11,55,56^

Nevertheless, we did not observe significant differences when comparing the DLB patients and their subgroups (MCI-DLB, d-DLB) with the MCI-AD and d-AD subgroups. However, this may be due to the limited sample size within the AD subgroups, reducing statistical power and obscuring potential trends.

Overall, we observed more pronounced global connectivity impairments in d-DLB patients than in AD patients, consistent with the notion that DLB is primarily characterized by functional rather than structural neurodegeneration. While widespread atrophy is a defining feature of AD, DLB typically shows relatively preserved brain structure,^40,42^ alongside alterations in the functioning of large-scale brain networks. Importantly, these impairments predominantly affected intra-network connectivity, indicating internal dysregulation within these core networks. This pattern stands in contrast to AD, where more extensive connectivity reductions are typically observed.^57^

Together, these observations support the view of DLB as a functionally driven disorder. Resting-state fMRI connectivity of large-scale brain networks like the SN, FPN or DMN could serve as markers to differentiate DLB from AD and other dementias, and provide insights into the mechanisms underlying the differential disease trajectories.

### Limitations

While our study provides comprehensive data on functional connectivity alterations in DLB, limitations warrant consideration. First, the relatively small sample size of the AD group may have reduced statistical power and limited generalizability. This could, for example, account for the unexpected absence of specific DMN alterations in the AD group when compared to the HC group. This issue was further exacerbated when subdividing the AD group into subgroups (11 MCI-AD and 14 d-AD) resulting in insufficient statistical power to detect significant correlations with a *p*_FDR_ < 0.05. Future studies should include larger AD cohorts to ensure sufficient statistical power for subgroup analyses and to validate these findings across disease stages.

Additionally, a longitudinal design in the fMRI analyses would help to clarify the temporal evolution of connectivity changes in DLB, providing insights into the trajectory of network dysfunction, helping to determine whether specific alterations precede clinical worsening or emerge because of disease progression.

Finally, although resting-state fMRI reveals valuable insights in baseline network organization, it lacks the specificity of task-based fMRI, which could better link connectivity changes to distinct cognitive or motor functions. Moreover, exploring additional analysis modalities, such as dynamic functional connectivity, may further elucidate the network alterations underpinning the complex symptomatology of DLB.^58,59^

## Conclusion

SN and FPN disruptions play a central role in the progression of DLB towards dementia, with distinct connectivity patterns that clearly differentiate it from AD and HC. These alterations were linked to core DLB symptoms and cognitive decline, whereas in AD, cognitive decline was more closely associated with DMN impairments. This divergence suggests distinct network-level mechanisms underlying the two diseases. Future research should explore dynamic functional connectivity to better capture the temporal evolution of these disruptions and refine our understanding of the neural bases of DLB, AD and related disorders.

## Acknowledgements

The authors thank all the patients and control subjects for their participation in the study.

## Author contributions

**Vincent Gabriel:** conceptualization; formal analysis; methodology; validation; writing-original draft. **Elena Chabran:** conceptualization; methodology; supervision; validation; writing-review & editing. **Marion Sourty:** methodology; software; writing-review & editing. **Benjamin Cretin**: investigation; validation; writing-review & editing. **Nathalie Philippi:** investigation; validation; writing-review & editing. **Candice Muller:** investigation; validation; writing-review & editing. **Pierre Anthony:** investigation; validation; writing-review & editing. **Catherine Demuynck:** investigation; validation; writing-review & editing. **Paulo Loureiro de Sousa:** investigation; resources; writing-review & editing. **Anne Botzung:** data curation; project administration; writing-review & editing. **Léa Sanna:** data curation; project administration; writing-review & editing. **Olivier Bousiges:** conceptualization; methodology; supervision; validation; writing-review & editing. **Frédéric Blanc:** conceptualization; investigation; methodology; supervision; validation; writing-review & editing; funding acquisition.

## Ethical Considerations

This research was approved by the local ethical committee (“Comité de Protection des

Personnes Strasbourg Est IV” [CPP-EST-IV]).

## Consent to Participate

All participants provided written informed consent to participate.

## Data Availability Statement

The data supporting the findings of this study are available on request from the corresponding author. The data are not publicly available due to privacy or ethical restrictions.

