## Supplemental Table 1 for "Network Signatures of Disease Progression and Core Symptoms in Dementia with Lewy Bodies Distinct from Alzheimer’s Disease"

**Supplementary Table 1. Sociodemographic and Clinical Characteristics of the MCI-DLB, d-DLB, MCI-AD, d-AD subgroups and the HC group.**

| Characteristic | | DLB | | AD | | | | |
|---|---|---|---|---|---|---|---|---|
| | | **MCI-DLB** *n* = 63 | **d-DLB** *n* = 27 | **MCI-AD** *n* = 11 | **d-AD** *n* = 14 | **HC** *n* = 34 | **Test statistics** [d], ***p*** | **Post-hoc analyses** |
| **Age (years)** | | 67.3 ± 9.0 | 74.8 ± 8.7 | 74.0 ± 6.9 | 73.1 ± 9.4 | 68.1 ± 8.1 | F = 4.87 $p < 0.001^{***}$ | d-DLB > MCI-DLB |
| **Gender (F/M)** [a] | | 38/25 | 15/12 | 6/5 | 9/5 | 21/13 | $\chi^2 = 3.22$ $p = 0.522$ | / |
| **MMSE score** | | 27.3 ± 2.0 | 21.8 ± 4.2 | 26.5 ± 2.0 | 22.0 ± 2.4 | 28.9 ± 1.0 | H = 83.01 $p < 0.001^{***}$ | HC > MCI-DLB, MCI-AD, d-DLB, d-AD; MCI-DLB, MCI-AD > d-DLB, d-AD |
| **Socio-educational level** [b] | | 12.3 ± 3.9 | 10.4 ± 4.6 | 12.6 ± 3.1 | 11.1 ± 3.0 | 13.2 ± 2.8 | H = 7.83 $p = 0.098$ | / |
| **IADL** | | 3.7 ± 0.6 | 2.7 ± 1.4 | 3.3 ± 0.9 | 3.2 ± 0.9 | 4.0 ± 0.0 (1 ND) | H = 32.32 $p < 0.001^{***}$ | HC > d-DLB, d-AD; MCI-DLB > d-DLB |
| **Fluctuations** | | 1.8 ± 1.2 | 2.1 ± 1.1 | 0.4 ± 1.0 | 0.6 ± 0.8 | 0.3 ± 0.5 | H = 57.06 $p < 0.001^{***}$ | MCI-DLB, d-DLB > HC, MCI-AD, d-AD |
| **Hallucinations** | | 1.8 ± 1.8 | 1.7 ± 2.1 | 0.3 ± 0.5 | 0.1 ± 0.4 | 0.2 ± 0.4 | H = 35.88 $p < 0.001^{***}$ | MCI-DLB, d-DLB > HC, MCI-AD, d-AD |
| **Parkinsonism** | **Akinesia** | 0.6 ± 0.6 | 1.3 ± 0.7 | 0.1 ± 0.3 | 0.0 ± 0.0 | 0.0 ± 0.2 | H = 67.85 $p < 0.001^{***}$ | MCI-DLB, d-DLB > HC, MCI-AD, d-AD; d-DLB > MCI-DLB |
| | **Rigidity** | 0.7 ± 0.6 | 1.2 ± 0.8 | 0.0 ± 0.0 | 0.1 ± 0.3 | 0.0 ± 0.2 | H = 60.01 $p < 0.001^{***}$ | MCI-DLB, d-DLB > HC, MCI-AD, d-AD; d-DLB > MCI-DLB |
| | **Tremors at rest** [c] | 31.75% | 29.63% | 10.00% | 0.00 % | 6.90% | $\chi^2 = 13.18$ $p < 0.01^{**}$ | MCI-DLB, d-DLB > HC, d-AD |
| **RBD** | | 1.1 ± 0.9 (2 ND) | 1.3 ± 0.8 | 0.3 ± 0.7 | 0.1 ± 0.4 | 0.4 ± 0.6 | H = 33.71 $p < 0.001^{***}$ | MCI-DLB, d-DLB > HC, MCI-AD, d-AD |

All values are presented as means ± standard deviation unless otherwise stated.

[a] Sex-ratio (Female/Male). [b] Number of years of education. [c] Percentage of patients with deficits. [d] Statistical analysis performed using the one-way ANOVA (F) with Tukey's multiple comparison test; the Kruskal-Wallis with Dunn's multiple comparison post-hoc test (H); or the chi-square test ($\chi^2$) with Bonferroni correction for multiple comparisons.

DLB, dementia with Lewy bodies; AD, Alzheimer's disease; ; MCI-DLB, dementia with Lewy bodies in the mild cognitive impairment stage; MCI-AD, Alzheimer's disease in the mild cognitive impairment stage; d-DLB, dementia with Lewy bodies in the demented stage; d-AD, Alzheimer's disease in the demented stage; HC, healthy controls; MMSE, Mini-Mental State Examination; IADL, Instrumental Activities of Daily Living; RBD, REM sleep behavior disorder.
